# ARTEMIS integrates autoencoders and Schrödinger Bridges to predict continuous dynamics of gene expression, cell population and perturbation from time-series single-cell data

**DOI:** 10.1101/2025.01.23.634618

**Authors:** Sayali Anil Alatkar, Daifeng Wang

## Abstract

Cellular processes like development, differentiation, and disease progression are highly complex and dynamic (e.g., gene expression). These processes often undergo cell population changes driven by cell birth, proliferation, and death. Single-cell sequencing enables gene expression measurement at the cellular resolution, allowing us to decipher cellular and molecular dynamics underlying these processes. However, the high costs and destructive nature of sequencing restrict observations to snapshots of unaligned cells at discrete timepoints, limiting our understanding of these processes and complicating the reconstruction of cellular trajectories. To address this challenge, we propose ARTEMIS, a generative model integrating a variational autoencoder (VAE) with unbalanced Diffusion Schrödinger Bridge (uDSB) to model cellular processes by reconstructing cellular trajectories, reveal gene expression dynamics, and recover cell population changes. The VAE maps input time-series single-cell data to a continuous latent space, where trajectories are reconstructed by solving the Schrödinger bridge problem using forward-backward stochastic differential equations (SDEs). A drift function in the SDEs captures deterministic gene expression trends. An additional neural network estimates time-varying kill rates for single cells along trajectories, enabling recovery of cell population changes. Using three scRNA-seq datasets—pancreatic *β*-cell differentiation, zebrafish embryogenesis, and epithelial-mesenchymal transition (EMT) in cancer cells—we demonstrate that ARTEMIS: (i) outperforms state-of-art methods to predict held-out timepoints, (ii) recovers relative cell population changes over time, and (iii) identifies “drift” genes driving deterministic expression trends in cell trajectories. Furthermore, *in silico* perturbations show that these genes influence processes like EMT. The code for ARTEMIS: https://github.com/daifengwanglab/ARTEMIS.

## 1 Introduction

Cellular processes such as differentiation (e.g., stem cell differentiation into pancreatic *β*-cells [28]), development (e.g., embryogenesis in zebrafish [6]), and disease progression (e.g., epithelial-mesenchymal transition (EMT) in A549 lung cancer cells [3]) are dynamic and highly complex. The advent of single-cell sequencing technologies has enabled the capture of gene expression at cellular resolution. However, these experiments are often expensive and destructive, yielding only snapshots of unaligned cells at discrete timepoints. This limitation impedes the analysis of cellular processes, as continuous gene expression across timepoints is unavailable, and reconstructing dynamic trajectories without cell lineage tracking remains challenging. Moreover, cellular processes typically involve continuous population changes, due to cell birth, proliferation, and death (Figure 1a.). A model that reconstructs cellular trajectories from discrete timepoints while accounting for population changes and identifying key genes driving cellular processes is crucial (Figure 1b.).

**Figure 1.**
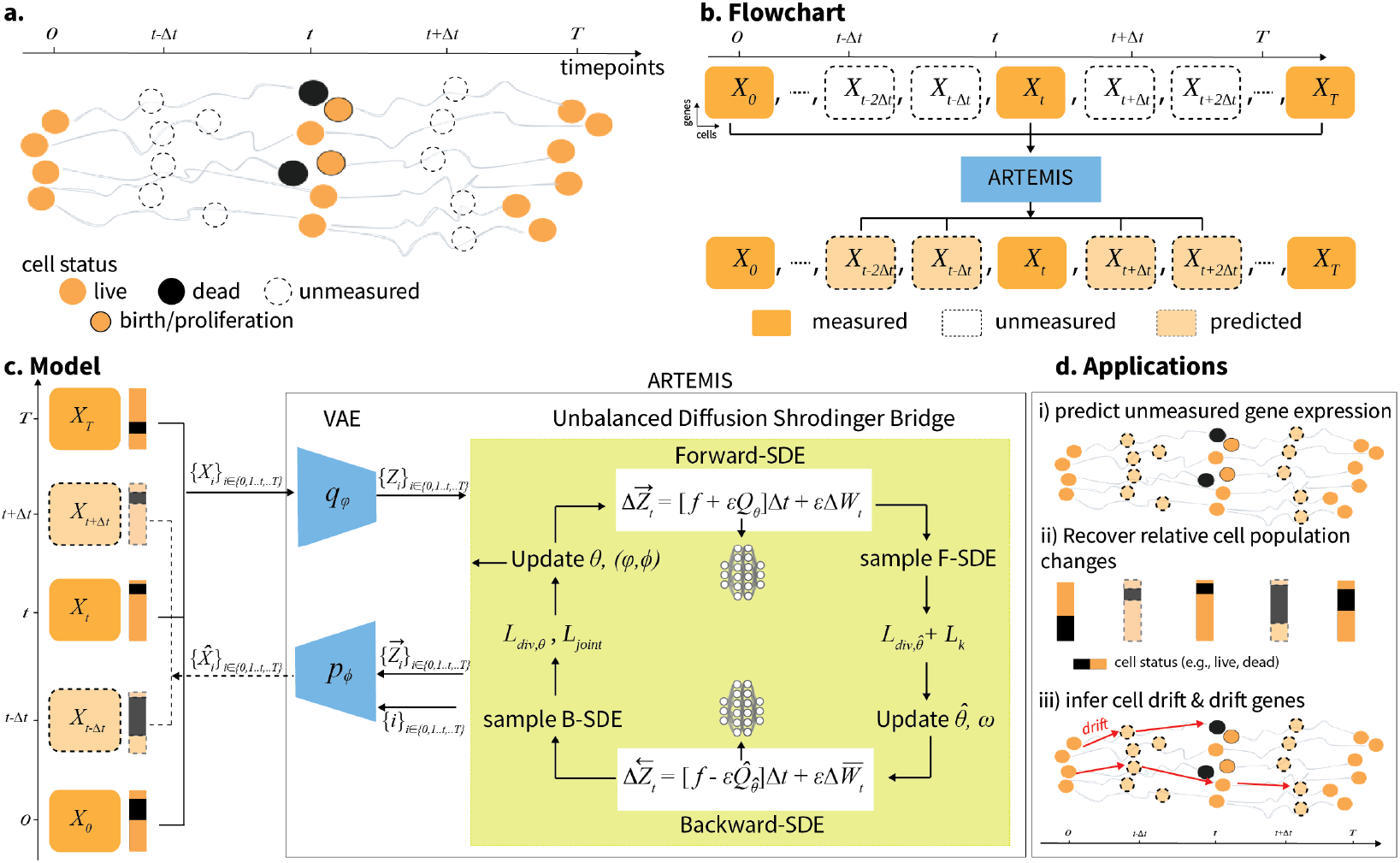
Model overview. a) Cellular processes are complex and dynamic, and undergo cell population changes driven by birth, proliferation and death with time, b) Single-cell sequencing provides snapshots of unaligned cells at discrete timepoins. To reconstruct cellular trajectories, we propose ARTEMIS, c) ARTEMIS leverages single-cell time-series gene expression data. It integrates and jointly trains a Variational Autoencoder (VAE) and unbalanced diffusion Schrödinger Bridge (uDSB) to learn a smooth latent space. The uDSB solves the SB problem using forward and backward SDEs, learning optimal backward and forward drifts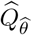, along with VAE parameters *φ,ϕ* by optimizing 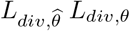 and, *L*_*joint*_, respectively. To further learn cell population changes, an additional loss *L*_*ω*_ is optimized, and d) ARTEMIS allows following downstream analysis i) predict gene expression for unmeasured timepoints, ii) recover relative cell population changes (infer cell status e.g., birth, proliferation, and, death) across timepoints, and, iii) learn cell drift and identifies drift genes.

Numerous methods have been developed for trajectory inference in single-cell gene expression data. Pseudotime analysis orders cells along trajectories based on gene expression similarity [9, 27, 31]. RNA velocity methods predict future transcriptional states by analyzing unspliced and spliced mRNA ratios in scRNA-seq data [8, 22]. However, both approaches rely on single snapshots, lacking temporal measurements, and thus limiting their ability to fully capture the dynamics of cellular processes.

Dynamic model-based approaches aim to address these limitations by learning continuous trajectories in real time. For example, PRESCIENT employs stochastic differential equations (SDEs) to model cellular differentiation, incorporating both deterministic and stochastic effects [32]. An SDE is formulated as d*x*_*t*_ = *b*(*x*)d*t* + *ε*d*W*_*t*_, where the drift term *b*(*x*) reflects deterministic trends derived from an energy potential’s gradient. PRESCIENT uses neural networks to learn this energy landscape. PI-SDE extends this approach with a physics-informed Hamilton-Jacobi (HJ) loss to improve training stability and identify least-action paths [15]. While these methods use growth rates derived from prior knowledge, they do not model population changes from cell birth, proliferation, or death events.

Optimal transport-based methods, like Waddington-OT, use unbalanced OT to infer probabilistic couplings between successive timepoints [24], but these approaches learn static, linear mappings. TrajectoryNet combines dynamic OT and continuous normalized flows, using neural ordinary differential equations (ODEs) to model cell population dynamics over time [26]. MIOFlow improves upon this by combining SDEs with a geodesic autoencoder, capturing non-linear latent spaces and preserving cellular variations [14]. However, these methods often struggle to generalize to unmeasured timepoints with distinct distributions due to fixed latent spaces. scNODE addresses this by combining a VAE with neural ODEs, incorporating dynamic regularization to enhance robustness against distributional shifts [33]. Nonetheless, ODE-based methods remain limited in capturing the stochasticity inherent in cellular processes.

Schrödinger bridges (SBs), a framework for dynamic entropy-regularized OT, have been applied to various fields including generative modelling [2, 5, 29], sampling [1, 13], and biological processes [12, 20]. SBs identify the most likely stochastic evolution between two probability distributions, given a prior or reference process (e.g., Brownian motion). Recent advances in diffusion models [25, 11] have led to algorithmic approximations called diffusion Schrödinger bridges (DSBs) [5]. By employing forward-backward SDE (FBSDE) theory, DSBs provide a computationally scalable approach to solve the SB problem [2]. Unbalanced DSBs (uDSBs) extend this framework by allowing unnormalized distributions, accommodating population changes such as cell death and proliferation [20]. However, SBs are prone to the curse of dimensionality making them difficult to scale to high-dimensional datasets like scRNA-seq [16].

Here, we present ARTEMIS (trAjectory infeRence wiTh unbalancEd dynaMic optImal tranSport), a generative model that integrates VAE with uDSBs to learn continuous gene expression dynamics and cell populations changes (Figure 1c). ARTEMIS first pre-trains a VAE to map scRNA-seq data into a low-dimensional latent space. The uDSB learns cellular trajectories by solving the SB problem through forward-backward SDEs, learning optimal forward-backward drift functions in this latent space. The VAE and uDSB are jointly trained to ensure a smooth latent space to predict cellular trajectories. Additionally, a neural network predicts time-varying kill rates, which are further used to infer cell statuses (e.g., birth, proliferation, death) along trajectories.

We benchmark ARTEMIS on three time-series scRNA-seq datasets and compare its performance to state-of-the-art methods, including PRESCIENT, MIOFlow, and scNODE, as well as using uDSB as a baseline. Our results demonstrate that ARTEMIS: i) accurately predicts single-cell gene expression at held-out timepoints, ii) recovers relative cell population changes over time, iii) learns a drift (*Q*) function in the SDE which captures deterministic trends in gene expression dynamics and identifies drift-genes along cellular trajectories. Furthermore, ARTEMIS enables the modeling of *in silico* perturbations introduced at intermediate timepoints (Figure 1d).

## 2 Materials and methods

### 2.1 Overview

ARTEMIS takes time-series scRNA-seq data for measured timepoints *t* ∈ {0, 1, 2, ‥, *T*} as input, and reconstructs the cellular trajectory from *t* = 0 to *t* = *T*. This includes gene expression for unmeasured timepoints 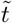, modeling relative cell population changes, and identifying genes associated with cell drift driving the trajectory. First, it pre-trains a VAE on single-cell gene expression 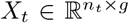 for measured timepoints *t*, given *n*_*t*_ cells and *g* genes by minimizing the *L*_*vae*_ loss (see 2.2). The VAE projects these cells to a *d*-dimensional (*d* ≪*g*) latent space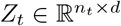. Then, the VAE and uDSB are jointly trained to model evolution of cellular trajectories from *t* = 0 to *t* = *T*. The uDSB training proceeds iteratively, alternating between forward and backward passes, with the VAE training integrated into the forward pass. In the forward pass, an optimal forward drift *Q*^*θ*^ is approximated by simulating a backward-SDE 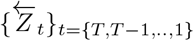, where 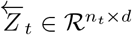 and minimizing the loss *L*_*div,θ*_ (see 2.3). The VAE is jointly trained in this forward pass by minimizing latent and reconstruction losses *L*_*joint*_ (see 2.4). Then, in the backward pass, an optimal backward drift 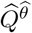is approximated by simulating a forward-SDE 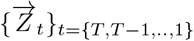, where 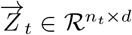 and minimizing the loss 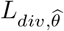 (see 2.3). Additionally, to predict cell population changes, another neural network *K*_*ω*_ is learned which determines how cell statuses (e.g., live, dead) (see 2.2) change. Once trained, ARTEMIS learns the optimal parameters for the VAE (*φ, ϕ*) and uDSB 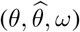. The pseudo-code to train ARTEMIS is summarized in Algorithm S1.

### 2.2 Learning interpretable latent space with variational autoencoders

The Variational Autoencoder (VAE) [17] has been one of the most popular generative models. The basic idea of VAE can be summarized as follows: (1) VAE encodes the input data samples into a latent variable as its distribution of representation via a probabilistic encoder, which is parameterised by a neural network. (2) then adopts the decoder to reconstruct the original input data based on the samples from the latent variable. Here, we pre-trained a VAE on scRNA-seq data for observed timepoints, where an encoder module *q*(·; *φ*) : ℝ ^*g*+1^ → ℝ ^*d*^ maps cells concatenated with sinusoidal encoded time (*X*_*t*_||*t*) to a latent space parameterized by a Normal distribution 𝒩 (*μ, σ*^2^), and a decoder *p*(·; *ϕ*) : ℝ ^*d*+1^ ← ℝ ^*g*^ maps (*Z*_*t*_||*t*) back to gene expression space:

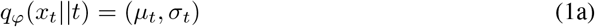

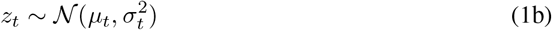

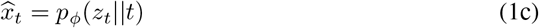

The encoder and decoder networks are parameterized by *φ* and *ϕ*, respectively. Then, the loss function *L*_*vae*_ minimized includes i) mean squared error (MSE) between the input and reconstructed scRNA-seq data, and, ii) the Kulback-Leibler (KL) divergence between the encoder output and a standard normal prior:

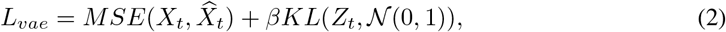

where *β* is a scaling factor for the KL-divergence term to ensure that the model learns robust and interpretable latent representations [10].

### 2.3 Modeling latent time-series with Schrödinger bridges

The Schrödinger bridge (SB) is the solution to an entropy-regularized optimal transport problem of finding the most likely evolution between two probability distributions. It seeks to find an optimal pair of forward-backward stochastic processes (SDEs) of the forms:

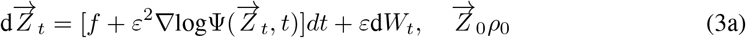

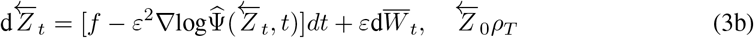

where (*ρ*_0_,*ρ*_*T*_) are the boundary distributions such that 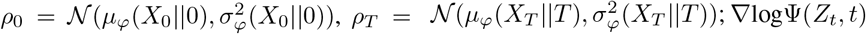 and 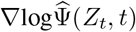 are the optimal forward and backward drifts, 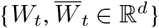 are standard Wiener processes and it’s time reversal, and, *f* and *ε >* 0 are base drift and diffusion coefficient, respectively. Both *f* and *ε* are constants known in prior. Moreover, the two SDEs in (3) can be thought of as *reversed* to each other. Now, suppose Ψ,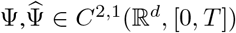 solve the following coupled PDEs,

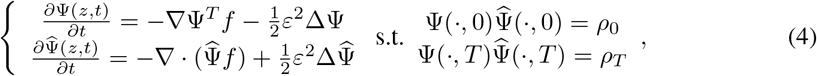

where ∇· is the divergence operator and Δ is the laplace operator. Then according to the SB theory, the solution to (4) can be expressed through the two coupled SDEs in (3)[18]. Here, the optimal forward (∇logΨ(*Z*_*t*_, *t*)) and backward 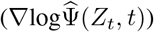 drifts are generally unknown and can be learned through neural networks parameterized by *θ*,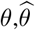, i.e.

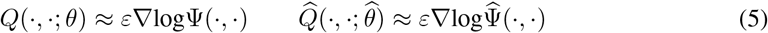

and is called the diffusion Schrödinger bridge (DSB). However, due to coupling constraints at the boundaries, solving (4) is a daunting task. Recently, [2] introduced a likelihood training framework grounded on forward-backward SDE (FB-SDE) theory which allows to construct likelihood objectives for training DSBs. Then the negative likelihood loss functions to solve for *θ* and 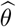 are:

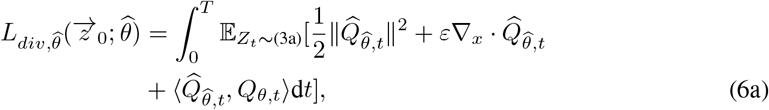

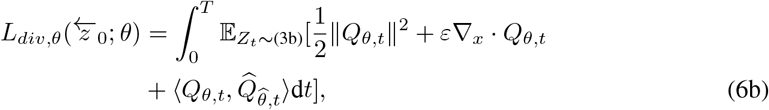

where ∇_*x*_· denotes the divergence operator with respect to the variable x: For any *υ* : ℝ^*d*^ → ℝ^*d*^, 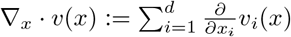. While DSBs assume that extreme/boundary marginals are normalized distributions, unbalanced DSBs (uDSBs) [20] relax this constraint by considering marginals with arbitrary mass i.e. by incorporating cell birth and death mechanisms. This is done by extending the state space ℝ^*d*^ to its *one-point compactification*, denoted as 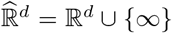, in which the added point ∞ serves as a “cemetery” or “coffin” state. This allows for jumps in processes, where new cells are born when state changes from ∞ → ℝ^*d*^ or existing cells are killed when ℝ^*d*^ → ∞. In this paper, we assume that in the absence of prior information about cell population changes, the number of cells (*n*_*t*_) collected at each timepoint represents the relative cell population. Also, we assume that any the variability introduced by sequencing technologies, such as incomplete sequencing of cells, is negligible when modeling the relative cell population changes over time. Then, in addition to estimating *θ* and 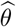, a posterior kill rate is learned using another neural network *K*(·; *ω*):

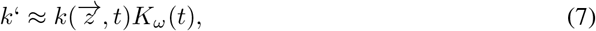

where *k*(*z, t*) *>* 0 is the prior kill rate. We follow uDSB to define the prior kill rate for a cell, which is defined as the ratio of the number of features deviating by more than two standard deviations (from the mean of expression of cells from the next measured timepoint) to the total number of features for that cell. If fewer than 20% of the features deviate, the prior kill rate is set to 0. The loss function to optimize *ω* is given by:

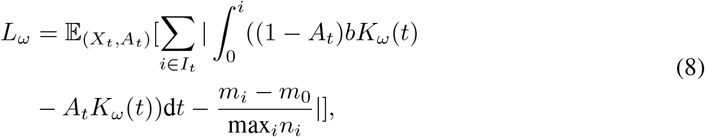

where 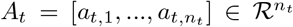 where *a*_*t,i*_, ∈ {0,1} for all *i* = 1, …, *n*_*t*_ such that *a*_*t,i*_ = 1 indicates live and *a*_*t,i*_ = 0 indicates dead cell status for some cell *i, b* is the birth rate (i.e. negative death rate) and *I*_*t*_ is the set of intermediate timepoints. The first three terms ensure that the change in mass predicted by uDSBs in each time interval [0,*i*] matches the empirical change (*n*_*i*_ − *n*_0_) for observed timepoints *t* ∈ *I*_*t*_. Additionally, we use the Euler-Maruyama discretization [20, 2] to approximate the SDEs from (3):

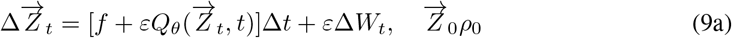

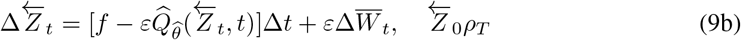

where the interval [0, *T*] is discretized into 100 steps. Then the descritized loss *L*_*k*_(*ω*) is:

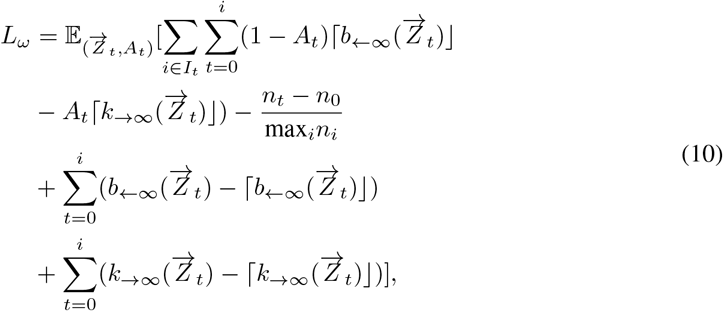

The last two regularization terms are added to penalize transition probabilities *k* _→_ ∞ and q _←_ ∞ greater than 1, where

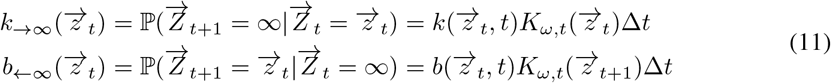

The discretized divergence losses (6a), (6b) become:

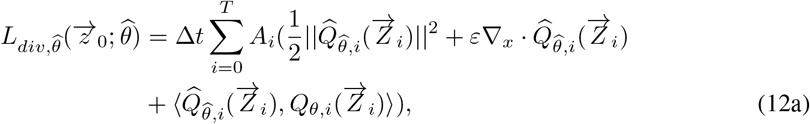

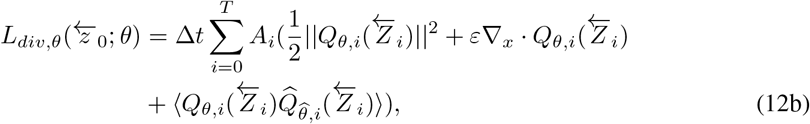

where 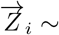 (9a) and 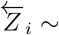 (9b). The sampling procedure for forward and backward SDEs with birth and death mechanisms is summarized in Algorithm S2 where the predicted kill rate *r*’ determines if cells are born,proliferate or die.

To learn optimal forward and backward drifts 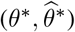, the iterative proportional fitting (IPF) algorithm [7, 23], which is the dynamic version of the sinkhorn algorithm, [4] has been used widely. It works by alternating between two steps, (i) a forward pass that adjusts the predicted distribution of a forward SDE ((9a)) to match the terminal distribution i.e. 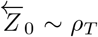, and (ii) a backward pass that adjusts the predicted distribution of a backward SDE ((9b)) to match the initial distribution i.e. 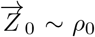. This process is repeated iteratively, with each iteration bringing the predicted distributions closer to satisfying marginal distributions. The training details are summarized in Supplementary Note S1 and Algorithm S1. For more details about uDSBs, we refer readers to [20].

### 2.4 Joint optimization of VAE and Schrödinger bridges for smooth time-series dynamics in latent space

Following pre-training of the VAE on gene expression data from measured timepoints, we jointly train the VAE and the uDSB to learn a smooth latent space for interpolation to unmeasured intermediate timepoints. As detailed in 2.4, the uDSB is trained iteratively through forward and backward passes to approximate optimal drifts while adhering to marginal distributions. However, this training process overlooks cellular profiles at intermediate timepoints. To address this, we introduce a latent loss term into the SB training, penalizing discrepancies between gene expression predictions in the latent space generated by the VAE and the uDSB. Additionally, the VAE is jointly trained with the forward pass, incorporating two additional loss functions into *L*_*div,θ*_(*z*_*T*_ ; *θ*) to minimize the distance: i) between VAE-encoded latents *Z*_*φ,t*_ and uDSB-predicted latents *Z*_*t*_, and ii) between the ground truth and VAE-reconstructed gene expression:

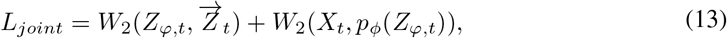

where *W*_2_(*μ, ν*) calculates the 2-Wasserstein distance between empirical distributions *μ* and *ν*.

### 2.5 Model Outputs

To reconstruct a complete trajectory from *t* = 0 to *t* = *T, X*_0_ is input to the trained model, which is mapped to the latent space by the encoder *q*_*φ*_. Then a forward trajectory 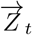 is sampled using (9a) upto *t* = *T*. Then 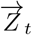 is mapped back to the gene expression using the decoder *p*_*ϕ*_ computing expression for measured(*t*) and unmeasured 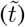 timepoints: 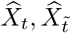. In addition to expression, ARTEMIS also outputs the cell status 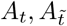 for cells in the reconstructed trajectory.

### 2.6 Datasets and preprocessing

We benchmarked ARTEMIS on three time-series scRNA-seq datasets (Table 1). The pancreatic dataset spans Days 0 to 7, capturing stage 5 human in vitro pancreatic *β*-cell differentiation [28]. The zebrafish dataset covers embryogenesis across twelve stages, measured in hours post-fertilization (hpf) [6]. Lastly, the EMT dataset involves an A549 lung cancer cell line treated with TGFB1 to induce epithelial-to-mesenchymal transition (EMT), sampled at five timepoints from 8 hours to 1 week post-treatment [3]. For all datasets, counts were normalized using depth scaling, ensuring that the total counts for each cell across all genes were consistent. This was followed by log1p normalization (log-transformation after adding one) to stabilize variance. These steps prevented data leakage between training and held-out data. Next, we identified 2,000 highly variable genes (HVGs) from the training data and applied this selection to the held-out samples. Preprocessing was performed using the *Scanpy* package [30]. For the EMT dataset, we additionally applied z-score normalization using the *scikit-learn* package [21], as we perform perturbation analysis on this data. To simplify computations, we relabeled timepoints as consecutive integers starting from 0. For the pancreatic dataset, we filtered out genes correlated with *TOP2A* (r *>* 0.15) following Yeo et al. (2021) before training resulting in 1922 HVG genes.

**Table 1:** Three scRNA-seq datasets using for benchmarking in this paper.

| Dataset | #cells | #timepoints | Source |
| --- | --- | --- | --- |
| Pancreatic β-cell differentiation (Pancreatic) | 51,274 | 8 | GEO (GSE114412) [28] |
| Zebrafish embryogenesis (Zebrafish) | 38,731 | 12 | Broad Single-Cell Portal SCP162 [6] |
| Epithelial-to-mesenchymal transition (EMT) | 3,133 | 5 | GEO (GSE147405) [3] |

**Table 2:** Wasserstein distance between the predicted and held-out timepoints. Numbers in **bold** indicate best performance.

| Datasets | Pancreatic |  | Zebrafish |  |  | EMT |
| --- | --- | --- | --- | --- | --- | --- |
|  | t=3 | t=6 | t=5.3 | t=7.0 | t=9.0 | t=2 |
| ARTEMIS | 9.84±0.008 | **9.05±0.009** | 30.38±0.04 | **28.96±0.07** | **30.38±0.04** | **45.1±0.01** |
| uDSB | 11.209±0.011 | 10.81±0.009 | 39.98±0.18 | 42.21±0.15 | 43.23±0.05 | 47.26±0.12 |
| scNODE | **9.73±0.0003** | 9.22±0.0005 | **29.8±0.002** | 29.2±0.09 | 31.8±0.003 | 46.2±0.005 |
| MIOFlow | 10.46±0.012 | 10.47±0.016 | 31.2±0.03 | 31.19±0.04 | 34.42±0.04 | 45.4±0.006 |
| Prescient | 10.36±0.011 | 11.12±0.032 | 50±0.15 | 47.67±0.14 | 49.2±0.2 | 47.01±0.02 |
| Prescient (w. growth rates) | 10.55±0.03 | 12.16±0.07 | - | - | - | 48.83±0.01 |

## 3 Results

### 3.1 Human in vitro *β*-cell differentiation in pancreas

ARTEMIS was first applied to a pancreatic *β*-cell differentiation dataset with eight timepoints [28]. To evaluate ARTEMIS’s predictive performance, we withheld timepoints 3 and 6 during training for comparison across methods, as 3 central to differentiation and 6 near the terminal stage. ARTEMIS demonstrated superior accuracy in reconstructing both timepoints, with better generalization to the later timepoint (*t* = 6), as measured by average Wasserstein distance (Table 1). This is illustrated by a 2D UMAP visualization of the predicted gene expressions (Figure 2a., Supplementary Figure S1a‥

**Figure 2.**
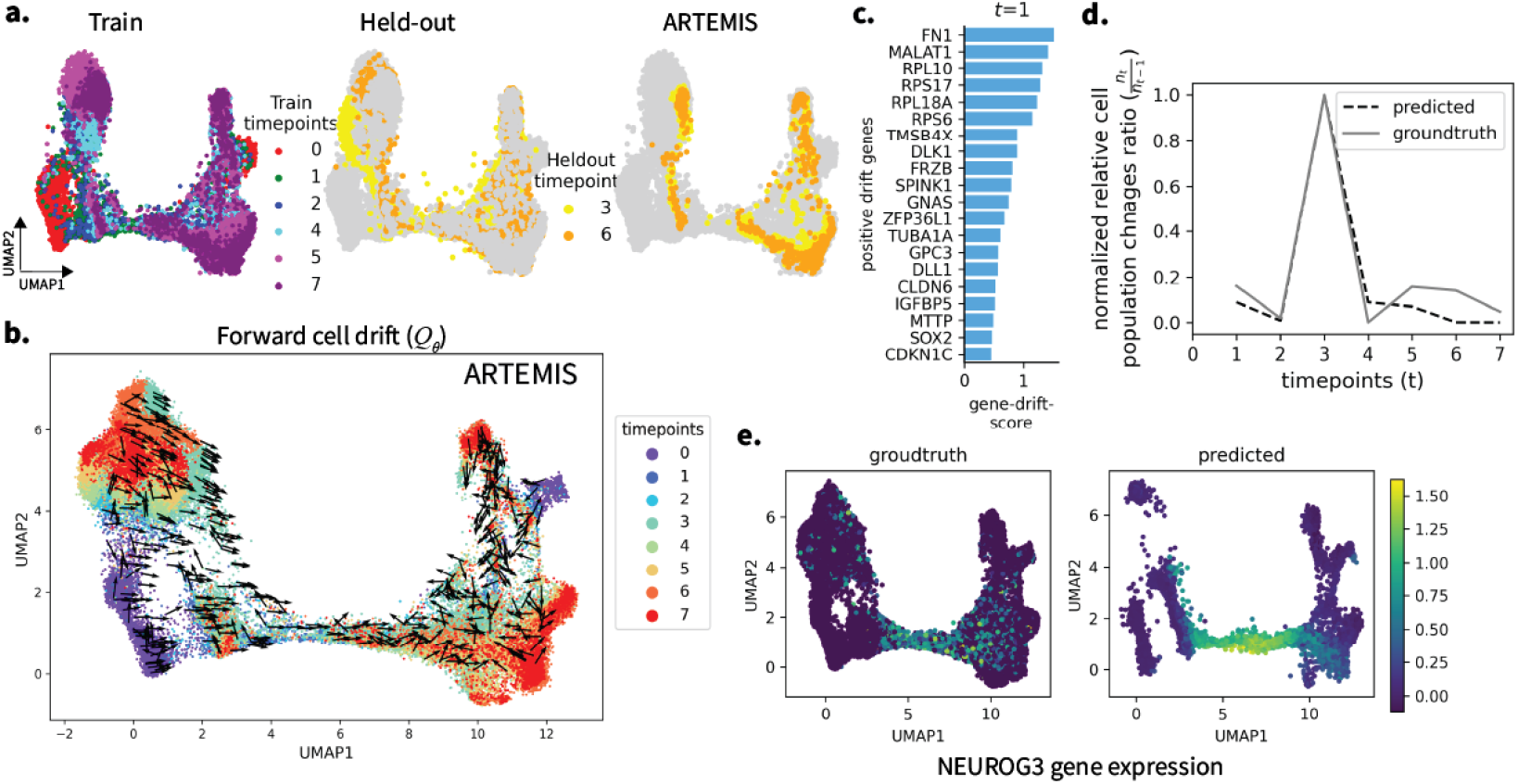
Application to pancreatic *β*-cell differentiation spanning eight days (0-7) a.) 2D UMAP to show ARTEMIS’s performance on held-out timepoints (3,6), b.) Visualization of the drift inferred by ARTEMIS trained on six timepoints, c.) Top 20 drift-genes identified for *t* = 1 from the forward drift *Q*^*θ*^, d.) Comparison of normalized ratios of relative cell population changes between ground truth and ARTEMIS-predicted cell statuses, e.) Groundtruth vs. predicted gene expression of transient TF *NEUROG3*

To further analyze the temporal evolution of cellular trajectories, we visualized the forward drift (*Q*_*θ*_(·,·)) inferred by ARTEMIS, representing deterministic trends in gene expression dynamics. ARTEMIS accurately modeled the progression of cells from progenitor states toward terminal states, such as exocrine and neurog3_early populations (*t*=0, lower left in Figure 2b.), by aligning drift directions with biological expectations. While PRESCIENT also infers cellular drift, it often misaligned directions, particularly in regions of bifurcation (Supplementary Figure S1b.).

To identify the key genes driving these cellular transitions, we performed drift-gene analysis by projecting the latent forward drift (*Q*^*θ*^(·,·)) onto the gene expression space. Assuming genes with positive drift scores as drivers of cellular trajectories, we identified the top 20 drift-genes at each timepoint (Figure 2c., Supplementary Figure S1d., see Supplementary Notes S5). At *t* = 1, genes such as *MALAT1* (*p* < 2.9*e*^−23^), *FN1*(*p* < 1.7*e*^−34^), and *SOX2*(*p* < 2.*ye*^−2^) were among the top-ranked and also previously known markers of stage 5 progenitor cells ([28]).

We then evaluated ARTEMIS’s ability to recover cellular population changes between *t* = 0 and *t* = 7. Relative cell population change ratios for groundtruth were calculated as 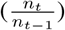 and normalized. Similarly, for ARTEMIS, these were given by 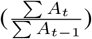 and normalized. We see that the trends captured by ARTEMIS closely matched groundtruth, suggesting that ARTEMIS can recover relative cell population changes for unmeasured timepoints (Figure 2c.) and inferred cell statuses capturing the increase and decrease in relative cell populations as birth and proliferation, and death, respectively (Supplementary Figure S1c.).

Additionally, ARTEMIS predicted the transient expression of *NEUROG3*, a critical transcription factor for endocrine induction and *β*-cell differentiation (Figure 2d.) [28] accurately. These results demonstrate ARTEMIS’s ability to integrate gene expression and population dynamics while identifying key genes driving cellular transitions.

### 3.2 Developmental stages underlying zebrafish embryogenesis

We next applied ARTEMIS to a zebrafish embryogenesis dataset spanning twelve developmental stages [6]. To evaluate prediction accuracy, three timepoints (*t* = 4, 6, 8) were held out during training as previously benchmarked by scNODE. ARTEMIS outperformed other methods in reconstructing these timepoints, with better generalization to later stages (Table 1), further illustrated by 2D UMAP visualizations of the predicted gene expressions (Figure 3a., Supplementary Figure S2a.).

**Figure 3.**
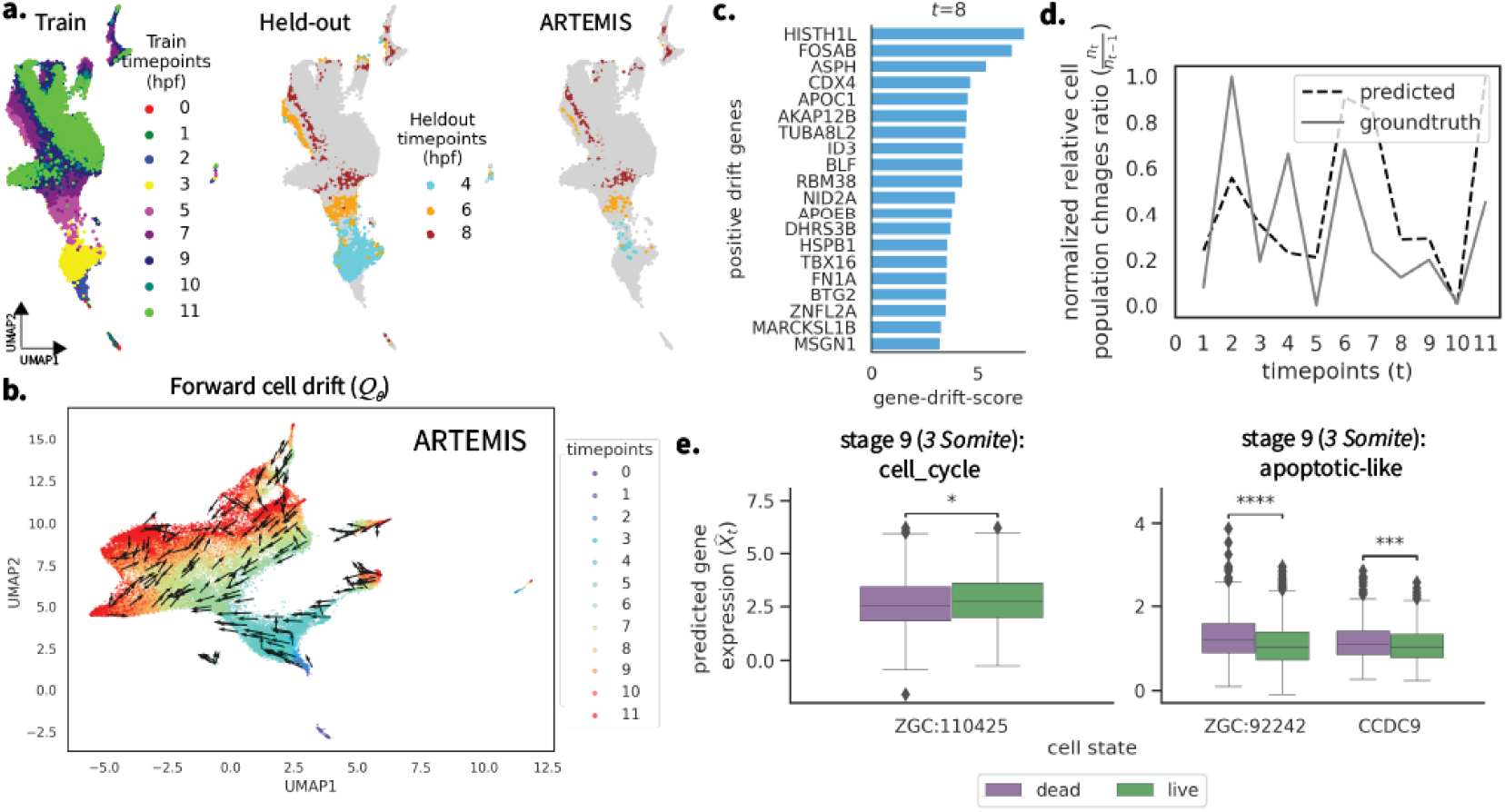
Application to zebrafish embryogenesis data across twelve stages (i.e. hours post fertilization (hpf)) a.) 2D UMAP to show ARTEMIS’s performance on held-out timepoints (4,6,8), b.) Visualization of the drift inferred by ARTEMIS trained on nine timepoints, c.) Top 20 drift-genes identified for *t* = 8, d.) Comparison of normalized ratios of relative cell population changes between ground truth and ARTEMIS-predicted cell statuses as live, e.)Boxplots showing gene expression of DE genes between cells predicted as live and dead by ARTEMIS during the interval *t* = 9 to *t* = 10.

To analyze developmental trajectories, we visualized the forward drift inferred by ARTEMIS and compared to PRESCIENT. ARTEMIS successfully modeled the deterministic evolution of cells through distinct developmental phases, aligning the drift consistently toward later stages of embryogenesis (Figure 3b., PRSCIENT: Supplementary Figure S2b.). Drift-gene analysis highlighted key regulators of zebrafish development at various stages (Figure 3c., Supplementary Figure S3d.). For example, at *t* = 8, top drift genes identified included *CDX4*(*p* < 2.9*e*^−6^), *APOC1*(*p* < 2.6*e*^−9^), and *TBX16*(*p* < 2.5*e*^−17^), known for their roles in Talibud formation. Similarly, genes such as *ID3* (*p* < 8.9*e*^−25^) and *FN1A*(*p* < 2.8*e*^−60^) are known for their role in Prechordal Plate lineage, and *ASPH* (*p* < 1.7*e*^−28^) and *APOEB*(*p* < 7.5*e*^−12^) in the Endoderm lineage. ([6]).

ARTEMIS also recovered cellular population changes across *t* = 0 to *t* = 11, closely matching the ground truth for relative cell population changes (Figure 3d.) and inferred cell statuses (Supplementary Figure S3c.). We also conducted DE analysis during the interval *t* = 9 to *t* = 10, where ARTEMIS predicted enough number of cells as dead. Here, ARTEMIS identified genes associated with the cell cycle, such as *ZGC:110425*(*p* < 0.05), as enriched in cells predicted live, while apoptotic-like genes, including *ZGC:92242*(*p* < 1*e*^−4^) and *CCDC9*(*p* < 1*e*^−3^), exhibited higher expression in cells predicted dead (Figure 3e.)[6]. This shows that ARTEMIS provides meaningful predictions of cell statuses, linking gene expression patterns to cellular states such as proliferation and apoptosis during key developmental dynamics.

### 3.3 Epithelial-to-mesenchymal transition in TFGB1-induced A549 lung cancer cells

Finally, we applied ARTEMIS to an Epithelial-to-mesenchymal (EMT) dataset of A549 lung cancer cells treated with TGFB1, spanning five timepoints. To evaluate predictive performance, *t* = 2 was held out during training as it is the central timepoint in the measured EMT process. ARTEMIS achieved the lowest average Wasserstein distance, outperforming other methods in reconstructing the held-out timepoint (Table 1), illustrated using a 2D UMAP visualization (Figure 4a., Supplementary Figure S3a.).

**Figure 4.**
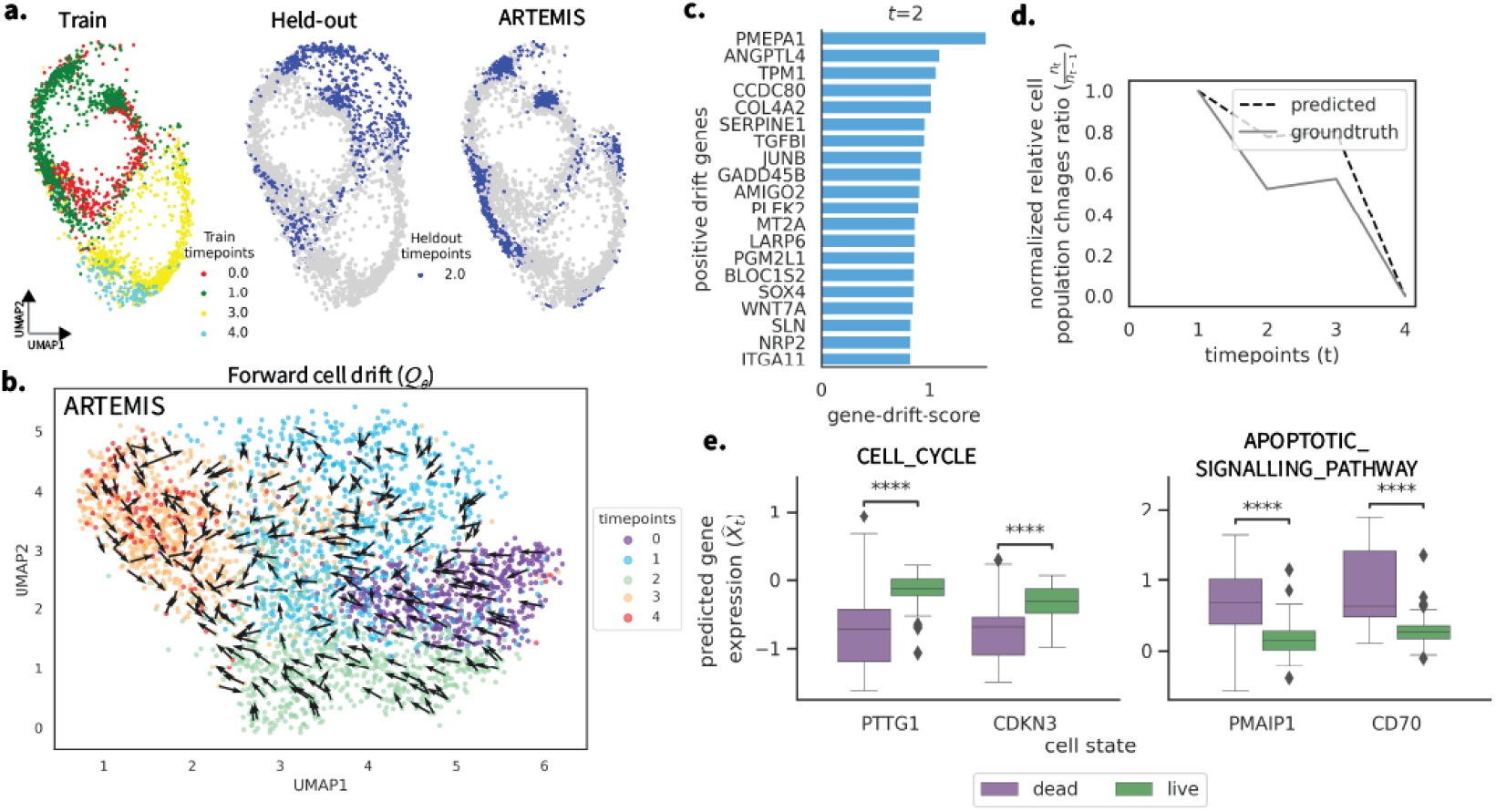
Application to A549 lung cancer cells undergoing TGFB1-induced EMT spanning five timepoints, a.) 2D UMAP to show ARTEMIS’s performance on held-out timepoint (4), b.) Visualization of the drift inferred by ARTEMIS trained on four timepoints, c.) Top 20 drift-genes identified for *t* = 2, d.) Comparison of normalized ratios of relative cell population changes between ground truth and ARTEMIS-predicted cell statuses as live, e.) Boxplots showing gene expression of DE genes between cells predicted as live and dead by ARTEMIS during the interval *t* = 3 to *t* = 4.

The forward drift inferred by ARTEMIS successfully captured the progression of cells toward later timepoints, modeling the deterministic trends associated with EMT (Figure 4b.). In contrast, PRESCIENT struggled to align directions accurately in complex regions (Supplemntary Figure S3b.). Drift-gene analysis identified key regulators of EMT at each timepoint (Figure 4c., Supplementary Figure S3d.). At *t* = 2, genes such as *COL42* (*p* < 5.7*e*^−10^), *PMEPA1* (*p* < 7.5*e*^−9^), *SERPINE1*(*p* < 6.2*e*^−4^), and *TPM1*(*p* < 1.9*e*^−15^) that were identified as top drift genes, are also known as EMT hallmark genes [19].

ARTEMIS also captured relative cell population changes across *t* = 0 to *t* = 4, closely matching the ground truth trends (Figure 4c.) and inferred cell statuses (Supplemetary Note S3c.). Differential expression (DE) analysis during *t* = 3 to *t* = 4, where most cells were predicted as dead, revealed distinct patterns. Genes previously associated with the GOBP CELL_CYLCE pathway, such as *PTTG1*(*p* < 1*e*^−4^) and *CDKN3*(*p* < 1*e*^−4^), were enriched in cells predicted as liv, while genes previously known in the GOBP APOPTOTIC_SIGNALLING_PATHWAY, including *PMAIP1*(*p* < 1*e*^−4^) and *CD70*(*p* < 1*e*^−4^), exhibited higher expression in cells predicted as dead (Figure 4e.) [19].

To further validate our findings and explore the role of identified drift genes *TPM1* and AMIGO2, we introduced *in silico* perturbations on cells at timepoint *t* = 2 by introducing varying levels of overexpression (5, 10, 15, 20, 25) or underexpression (25, 20, 15, 10, 5) of the drift genes (Figure 5, see Supplementary Note S6). The trained ARTEMIS model was initialized by 2000 randomly sampled cells from *t* = 2, and allowed to reconstruct remaining trajectory up to terminal timepoint *t* = 4. At *t* = 2, model was initialized with unperturbed cells and cells with perturbations. This was repeated for 10 trials. An MLP classifier, trained on ground truth latents inferred by the VAE, was used to classify the cells from the predicted trajectory into five timepoints. For each timepoint, a two-sided t-test (*p* < 0.05) compared the distribution of cells between perturbed and unperturbed groups. When underexpressed, there was an increase in cells from earlier timepoints (*t* = 0, 1) and a decrease in cells from later timepoints in perturbed trajectories, compared to the unperturbed trajectories (*p* < 0.001, Figure 5a). Conversely, overexpression resulted in increased cell populations at later timepoints (*t* = 3, 4) and a decrease at earlier timepoints (*p* < 0.05) in perturbed trajectories (Figure 5b). Similar patterns were observed across other perturbation levels (Supplementary Figure S4). These findings suggest that the drift genes identified by ARTEMIS not only reflect deterministic trends underlying EMT but can also influence cellular trajectories when perturbed.

**Figure 5.**
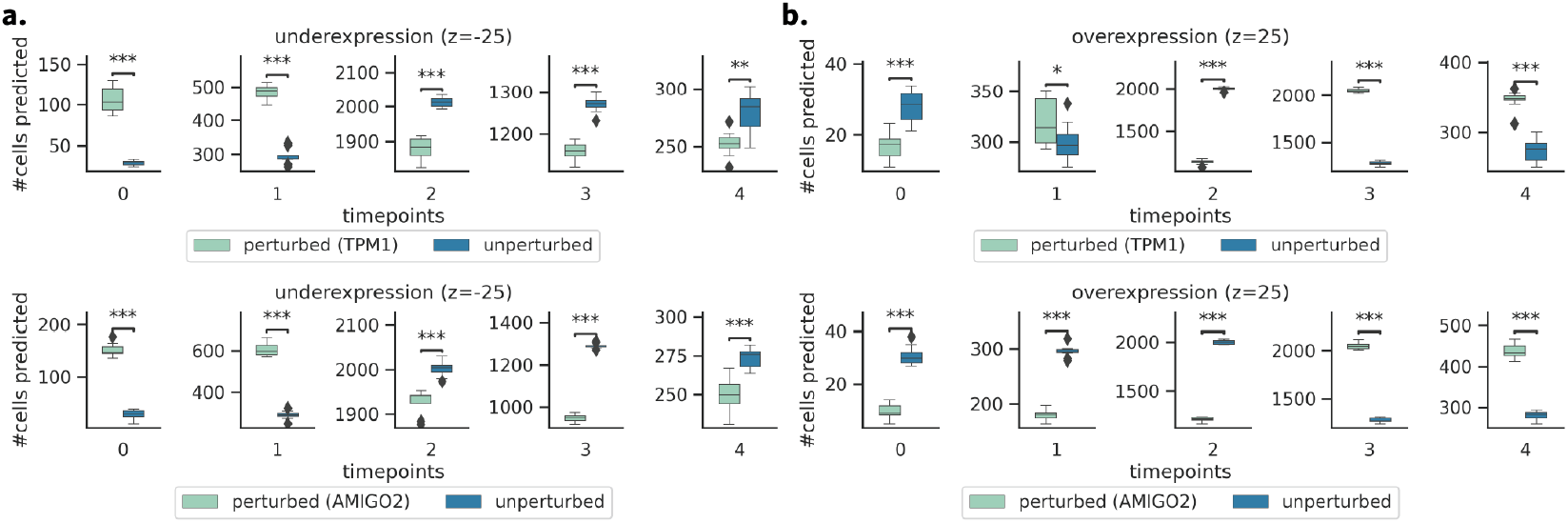
Perturbation analysis was conducted on A549 lung cancer data undergoing TGFB1-induced EMT. Boxplots illustrate the number of cells assigned to specific timepoints. Trained ARTEMIS model was initialized with 2000 cells randomly sampled from *t* = 2 either unperturbed or after introducing perturbations of TPM1 and AMIGO2, and simulate the trajectory until terminal timepoint. An MLP classifier assigned cells to specific timepoints. The number of cells asigned to each timepoint from perturbed and unperturbed trajectories were compared using a two-sided t-test at p < 0.05 (see Supplementary Note S6). (a) shows results for underexpressed genes with a perturbation value of -25, and (b) shows results for overexpressed genes with a perturbation value of 25

## 4 Discussion

Here we propose ARTEMIS, a generative model to reconstruct cellular trajectories, recover cell population changes, and learn cellular drift and drift-genes driving these trajectories.

ARTEMIS addresses challenges related to scalability and the curse of dimensionality often encountered in optimal transport problems [16] by leveraging a VAE to learn interpretable, low-dimensional representations of single-cell gene expression data, continuously optimized during training.

The joint training of complex models like VAE and uDSB introduces additional complexity, making hyperparameter tuning a non-trivial task. To assist users, we evaluate ARTEMIS across various hyperparameter settings (see Supplementary Note S4, Supplementary Figure S4.) to provide heuristic guidance on selecting appropriate configurations.

Predicting cell states (e.g., birth, proliferation, or death) at unmeasured time points remains challenging due to the complexity and rapid evolution of cellular processes. While ARTEMIS can recover relative cell population changes and predict cell states at these time points, incorporating prior knowledge—such as prioritizing genes associated with pathways like the cell cycle or apoptosis—during training could enhance its ability to better align with biological processes. Future work will focus on integrating such domain knowledge to improve biological fidelity.

We also aim to expand ARTEMIS by incorporating additional modalities, such as chromatin accessibility to gain insights into regulatory mechanisms and improve the prediction of biologically relevant trajectories.

## 7 Data availability

All the datasets used for analysis in this paper are publicly available. The pancreatic data, can be downloaded from GEO (GSE114412)[28]. The raw zebrafish data can be downloaded from https://figshare.com/articles/dataset/Raw_and_processed_data_of_three_scRNA-seq_datasets_/25601610/1?file=45647244[33]. It can also be downloaded from Broad Single Cell Portal with identifier SCP126 [6]. The TGFB1-induced EMT from A549 lung cancer cell data can be downloaded from https://github.com/dpcook/emt_dynamics [3].

