## Supplementary materials for "ARTEMIS integrates autoencoders and Schrödinger Bridges to predict continuous dynamics of gene expression, cell population and perturbation from time-series single-cell data"

### Supplementary Notes

**Algorithm S1** ARTEMIS

**Input**: $\{X_t\}_{t\in\{0,1,..T\}}, t \in \{0, 1, ..T\}$, prior kill rate $k$, pretrain epochs $E_1$, number of iterations $E_2$, joint train epochs $E_3$, #SDEs to sample $E_4$

**Initialize**: $q_\varphi, p_\phi, Q_\theta, \widehat{Q}_{\widehat{\theta}}, r^\omega$

```
1:  Pre-train VAE:
2:  for i=1 to E_1 do
3:      q_φ(X_t||t) = (μ_t, σ_t), Z_t ~ N(μ_t, σ_t^2), X̂_t = p_φ(Z_t)
4:      Update φ, ϕ using ∇L_vae (Eq. 2)
5:  end for
6:  Jointly train VAE and uDSB:
7:  for j=1 to E_2 do
8:      for k=1 to E_3 do
9:          for l=1 to E_4 do
10:             (Z⃖_t, A_t) ← sample B-SDE(Q_θ, Q̂_θ̂, K_ω) (Eq. 9b)
11:             Update θ using ∇_θ L_{div,θ} (Eq. 12b)
12:             sample F-SDE Z⃖_t ~ ρ_0 (Eq. 9a)
13:             update θ, (φ, ϕ) using ∇_{θ,φ,ϕ} L_joint (Eq. 13)
14:         end for
15:         for l=1 to E_4 do
16:             (Z⃗_t, A_t) ← sample F-SDE(Q_θ, Q̂_θ̂, K_ω)(Eq. 9a)
17:             update θ̂, ω using ∇_θ̂ L_{div,θ̂} (Eq. 12a),∇_ω L_k (Eq. eq:10), respectively
18:         end for
19:     end for
20: end for
21: Outputs:
        φ, ϕ, θ, θ̂, ω
```

#### S1 Details of model implementation, training, and testing

ARTEMIS is implemented in JAX [1]. Training begins with VAE pre-training on single-cell gene expression data from observed timepoints. The ADAM optimizer with gradient clipping (threshold=1) and initial learning rate as 0.0001 was used and the VAE was trained for $E_1$ epochs with a default batch size of 64 (Algorithm S1, Steps 1-5).

Subsequently, the uDSB and VAE are jointly trained to learn a smooth latent space. The uDSB model comprises three networks, a.) forward drift $Q_\theta$, b.) backward drift $\widehat{Q}_{\widehat{\theta}}$, and c.) kill rate $K_\omega$.

The architecture of $Q_\theta$ and $\widehat{Q}_{\widehat{\theta}}$ includes:

i) `x_encoder`: 3-layer MLP taking $Z_t$ as input, ii) `t_encoder`: 3-layer MLP taking the sinusoidal embedding of time $t$ as input, and, iii) `decoder`: 2-layer MLP decoder combining ouputs from `x_encoder` and `t_encoder` to output optimal drift values ($Q_\theta$,$\widehat{Q}_{\widehat{\theta}}$).

The $K_\omega$ network is a 2-layer MLP taking a sinusoidal embedding of time $t$ and outputting a kill rate at time $t$. `SiLU` activation was used for $Q_\theta$ and $\widehat{Q}_{\widehat{\theta}}$; `Leaky-ReLU` for $K_\omega$. All networks used 16-dimensional time embeddings. The ADAM optimizer with gradient clipping was used, with initial learning rates of 0.0001 for all networks (needs to be tuned according to the dataset).

uDSB training is performed within the latent space from the pre-trained VAE. The uDSB training used a batch size of 512 over $E_2$ iterations, each iteration comprising

**Algorithm S2** SDE Sampling Procedure

**Input**: drift $f$, diffusion coefficient $\varepsilon$, killing rate $r$‘, initial cells $n_0$, final cells $n_T$

**procedure** F-SDE($Q_\theta, \widehat{Q}_{\widehat{\theta}}, K_\omega$)
  sample $\overrightarrow{Z}_0 \sim \rho_0$
  $A_0 \leftarrow 1$
  **for** step i=1 to T **do**
    $\overrightarrow{Z}_i = \overrightarrow{Z}_{i-1} + \Delta\overrightarrow{Z}_i$
    **if** $A_{i-1} = 1$ **then**
      $D \sim \text{Bernoulli}(1 - k‘(i)\Delta t)$
      $A_{i+1} \leftarrow D$
    **end if**
  **end for**
**end procedure**
**procedure** B-SDE($Q_\theta, \widehat{Q}_{\widehat{\theta}}, K_\omega$)
  sample $\overleftarrow{Z}_0 \sim \rho_T$
  $A_T \sim \text{Bernoulli}(\min(1, \frac{n_T}{n_0}))$
  **for** step i=T-1 to 0 **do**
    $\overleftarrow{Z}_i = \overleftarrow{Z}_{i-1} - \Delta\overleftarrow{Z}_i$
    **if** $A_{i+1} = 0$ **then**
      $D \sim \text{Bernoulli}(k‘(i)\Delta t)$
      $A_i \leftarrow D$
    **end if**
  **end for**
**end procedure**
**Outputs:**
  $(Z_t, A_t)_t$

$E_3$ epochs of forward and backward training. The uDSB model was trained using the Iterative Proportional Fitting algorithm (IPF), which iteratively solves the schrödinger bridge problem through forward and backward SDEs[2, 3, 4]. During each epoch, the forward and backward drifts, as well as the VAE parameters, are updated (Algorithm S1, Steps 6-20). Specifically, in each epoch, 10 SDEs were sampled to optimize forward and backward drifts each. The process is described below:

1. Forward optimization: This step minimizes the KL divergence with a fixed terminal condition (e.g., $\overleftarrow{Z}_t \sim p_T$). A backward SDE is sampled (Eq. 9b), and the divergence loss is calculated between the SDE predicted by the forward drift and the sampled backward SDE. The VAE params are optimized concurrently using the $L_{joint}$ loss (Eq. 13, Algorithm S1, Steps 9-14))

2. Backward optimization: This step minimizes the KL divergence with a fixed initial condition (e.g., $\overrightarrow{Z}_t \sim p_0$). A forward SDE is sampled (Eq. 9a) and the divergence loss is calculated between the SDE predicted by the backward drift and the sampled forward SDE (Algorithm S1, Steps 15-18)).

The number of discretization steps was set to 100 for the inteval [0,T], so $\Delta t$=0.01.

For trajectory inference, gene expression profiles at $t = 0$ are projected into the VAE latent space. Forward SDE sampling (Eq. 9a) with the learned drift $Q_\theta$ generates continuous latent variables, decoded back into the gene expression space to reconstruct cellular trajectories.

Prediction performance on held-out timepoints was evaluated by averaging the 2-Wasserstein distance between predicted and ground-truth gene expression profiles, computed over five forward trajectory samples. The `OTT` library [5] was used to compute distances.

We performed all training and benchmakring on Linux Ubuntu machine with 256 BG RAM and NVIDIA RTX A6000 GPU with 48 GB RAM. The code has also been tested on Linux Ubuntu machine with only CPU.

#### S2 Baseline Methods

We compare ARTEMIS's performance with the following baseline methods:

- PRESCIENT [6]: PRESCIENT (Potential eneRgy undErlying Single Cell gradIENTs) is a generative modeling framework designed to learn differentiation landscapes from time-series scRNA-seq data. It models how cells evolve stochastically and in physical time, using a diffusion-based approach to recover a global potential function. To handle large scRNA-seq datasets, PRESCIENT models are fit on PCA projections of scaled gene expression data. The potential function is parameterized by a neural network, to allow flexible and complex landscape modeling. PRESCIENT allows for using prior knowledge of cell growth in the modeling. However, such information is not always available and we included evaluations with growth rates information when available. We used Python codes on Github (`https://github.com/gifford-lab/prescient-analysis`) to run PRESCIENT.
- MIOFlow [7]: MIOFlow (Manifold Interpolating Optimal-Transport Flow) is a computational method for modeling stochastic, continuous population dynamics from snapshots of time-series data. It combines dynamic models, manifold learning, and optimal transport techniques to interpolate between static population snapshots. Using neural ordinary differential equations (Neural ODEs) and a geodesic autoencoder (GAE), MIOFlow ensures the flow aligns with the data's manifold geometry. By operating in the autoencoder's and penalizing transport with Wasserstein distance, complex diffusion processes in cellular dynamics. We use Python codes on GitHub (`https://github.com/KrishnaswamyLab/MIOFlow`) to run MIOFlow
- scNODE [8]: scNODE (single-cell Neural Ordinary Differential Equation) is a deep learning model that predicts and simulates single-cell gene expression at unobserved timepoints in temporal scRNA-seq data. It combines a variational autoencoder (VAE) to encode gene expression into a low-dimensional latent space with neural ordinary differential equations (ODEs) to model the temporal evolution of cells within this space. A dynamic regularization term aligns the latent dynamics with temporal data, reducing information loss between discrete timepoints and improving predictions. We use the Python codes on Github (`https://github.com/rsinghlab/scNODE`) to run scNODE.
- uDSB [9]: Unbalanced Diffusion Schrödinger Bridge (UDSB) is an extension of the Diffusion Schrödinger Bridge (DSB) framework that allows for modeling the temporal evolution of populations with changing mass over time. Unlike traditional DSBs, which assume conservation of mass and work with probability measures, UDSBs can handle marginals with arbitrary finite mass. uDSBs achieve this by incorporating stochastic differential equations with killing and birth terms, and by deriving their time reversals. We use the Python codes on Github (`https://github.com/matteopariset/unbalanced_sb`) to run scNODE.

#### S2 Hyperparameter Tuning

To benchmark methods evaluated in this paper, we selected parameters based on the average 2-Wasserstein distance using 3-fold cross validation. Here, we split the cells in each training timepoint into 3 sets, and perform cross-validation such that 2 sets are used for training, and the third for testing. To search for optimal hyperparameters, we used *wandb* [10].

For baseline uDSB, we first used PCA to project the gene expression to 50-dimensional space. For the networks $Q_\theta$ and $\widehat{Q}_{\widehat{\theta}}$, the `x_encoder` was 3-layer MLP with 300-dimension hidden layers, the `t_encoder` was 3-layer MLP with 32 dimension hidden layers, with takes 16-dimensional sinusoidal embedding of time $t$, and, the `decoder` was a 3-layer MLP decoder with 300 dimension hidden layers accepts concatenation of outputs from `x_encoder` and `t_encoder`. The $K_\omega$ network includes a 5-layer MLP with 64-dimension hidden layers. As activation functions, the `SiLU` for $Q_\theta$ and $\widehat{Q}_{\widehat{\theta}}$, and, `Leaky-ReLU` for $K_\omega$ networks. The ADAM optimizer wth gradient clipping was used, and initial learning rates for $Q_\theta$ and $\widehat{Q}_{\widehat{\theta}}$ were set to 0.001, and 0.01 for $K_\omega$. A batch size of 512 was used, and total training was conducted over 10 iterations, where each iteration included 10 epochs of forward and 10 epochs of backward training.

For PRESCIENT, we searched over the following hyperparameter spaces: latent dimension ∈ {10,50}, number of hidden layers ∈ {1,2,3}, sd ∈ [0.0,1.0], tau ∈ [0.0,0.1], gradient clipping ∈ [0.0,1.0]. The remaining hyperparameters were set to default. We estimated the growth rates for the EMT [11] and pancreatic [12] datasets using the mean of z-scores annotated to birth (KEGG_CELL_CYCLE) and death ()KEGG_APOPTOSIS) as suggested in [6].

For MIOFlow, we searched over the following hyperparameter spaces: gae embedded dim ∈ {10,50}, layers ∈ {[50,50],[16,32,16]}, $\lambda$ ∈ [1,40,100]. The remaining hyperparameters were set to default.

For scNODE, we searched over the following hyperparameter spaces: latent dimension (d) ∈ {10,50}, encoder network size ∈ {None, [d], [d,d]}, decoder network size ∈ {None, [d], [d,d]}, drift network size ∈ {None, [d], [d,d]}.

For ARTEMIS, we searched over the following hyperparameter spaces: latent dimension ∈ {10,50}, vae encoder hidden dimension ∈ {[512,256], [256,128]}, vae decoder hidden dimension ∈ {[512,256], [256,128]}, vae pre-train epochs ($E_1$) ∈ {50,100}, number of iterations ($E_2$) ∈ {2,4,6,8,10}, all learning rates ∈ {0.001,0.0001}, vae batch size ∈ {32, 64}, SDE sampling batch size ∈ {256,512}.

#### S4 Investigate ARTEMIS hyperparameters

We next investigate the hyperparameters tuned in ARTEMIS and their effects on prediction performance at unmeasured timepoints. Using the zebrafish dataset [13], we evaluate its performance on the previously defined task where three ($\tilde{t} = 4, 6, 8$) out of twelve timepoints were held out. We hope this analysis provides heuristic guidance to users to choose hyperparameters for training ARTEMIS. We vary the following hyperparameters:

1. base drift $f$ from {0,2,4,8,10,50,100} (Supplementary Figure S5a.)
2. Number of discretization steps from {12,50,75,100} (Supplementary Figure S5b.)
3. VAE latent dimension $d$ from [10,25,50,75,100,125,175,200] (Supplementary Figure S5c.)
4. SDE sampling batch size from {32,64,128,256,512} (Supplementary Figure S5d.)
5. Effect of using $L_{joint}$ loss terms, i.e., including either, both, or none of the terms (Supplementary Figure S5e.)

#### S5 Identify drift-genes

To bridge the latent forward drift dynamics with gene expression changes, we map the learned latent drift values to the gene expression space. Let $\overrightarrow{z}_{t,i} \in \mathcal{R}^d$ be a latent variable generated using the forward SDE (Eq. 9a) for a cell $i$ at time $t$. Let $\widehat{x}_{t,i} \in \mathcal{R}^g$ be the reconstructed gene expression for the cell $i$ using the decoder ($p_\phi$). To map the forward drift at $t$ to the gene expression space, we multiply the output of $Q^\theta(\overrightarrow{z}_{t,i}, t) \in \mathcal{R}^d$ with the jacobian of $p_\phi$ to compute the jacobian vector product (JVP) [14].

The jacobian $J \in \mathcal{R}^{g\times d}$ of $p_\phi$ is given by:

$$J_i = \begin{bmatrix} \frac{\partial \hat{x}_{t,i,(1)}}{\partial \overrightarrow{z}_{t,i,(1)}} & \frac{\partial \hat{x}_{t,i,(1)}}{\partial \overrightarrow{z}_{t,i,(2)}} & .. & \frac{\partial \hat{x}_{t,i,(1)}}{\partial \overrightarrow{z}_{t,i,(d)}} \\ . & . & . & . \\ . & . & . & . \\ \frac{\partial \hat{x}_{t,i,(g)}}{\partial \overrightarrow{z}_{t,i,(1)}} & \frac{\partial \hat{x}_{t,i,(g)}}{\partial \overrightarrow{z}_{t,i,(2)}} & .. & \frac{\partial \hat{x}_{t,i,(g)}}{\partial \overrightarrow{z}_{t,i,(d)}} \end{bmatrix}, \tag{1}$$

Then, the drift scores for cell $i$ can be calculated as:

$$\text{drift\_scores}_i = J_i Q_\theta(\overrightarrow{z}_{t,i}, t), \tag{2}$$

where $\text{gene\_drift\_scores}_i \in \mathcal{R}^g$. This is then averaged over all cells at time $t$ to get an average score for each gene:

$$\text{drift\_scores} = \frac{\sum_{i=1}^{n_t} J_i Q^\theta(\overrightarrow{z}_{t,i}, t)}{n}, \tag{3}$$

where $\text{drift\_scores} \in \mathcal{R}^g$ and $\text{drift\_scores}_j$ gives a gene-drift-score for gene $j$. We used the `jax.jvp()` function from JAX library for computing forward-mode Jacobian-vector product.

##### *Significance testing of drift-genes*

To evaluate the significance of the gene-drift scores, we randomized the positions of cells in the latent space and calculated their forward drifts. The drift values from the randomized cells, combined with the decoded gene expression of the original unshuffled cells, were used to establish a null distribution for gene-drift scores. A two-sided $t$-test was then performed to compare the observed gene-drift scores against the null distribution. Most drift genes identified by ARTEMIS were found to be significant, with p-values $< 0.05$.

#### S6 Perturbation analysis

For the EMT dataset [11], *in silico* perturbations were introduced by modifying the scaled normalized expression of target genes to z-score values: less than 0 for underexpression (knockdowns) and greater than 0 for overexpression. The resulting perturbed gene expression profile was then input to the trained ARTEMIS model, where it is was projected to the latent space to generate a forward SDE up to time $T$. Perturbations were introduced with magnitudes of -25,-20,-15,-10,10,15,20,25.

We conducted 10 trials, where 2,000 cells sampled from the timepoint of perturbation introduction were used to predict cellular trajectory up to time $T$ using the pre-trained ARTEMIS model. These simulations allowed us to examine the effects of perturbations within the latent space.

We trained a multilayer perceptron (MLP) classifier using the Python library scikit-learn [15] to predict timepoints based on latent cell representations. For a given trajectory resulting from perturbations introduced at time $t$, the predicted trajectory in the latent space was classified using the pre-trained MLP classifier.

Then, the number of predicted cells classified into each timepoint were compared between perturbed and unperturbed trajectories by performing a two-sided $t$-test test. We used this analysis to identify if the introduced perturbations for the drift-genes could alter/reverse the epithelial-to-mesenchymal transition by generating more cells corresponding to earlier or later timepoints (Figure 5, Supplementary Figure S4).

### Supplementary Figures

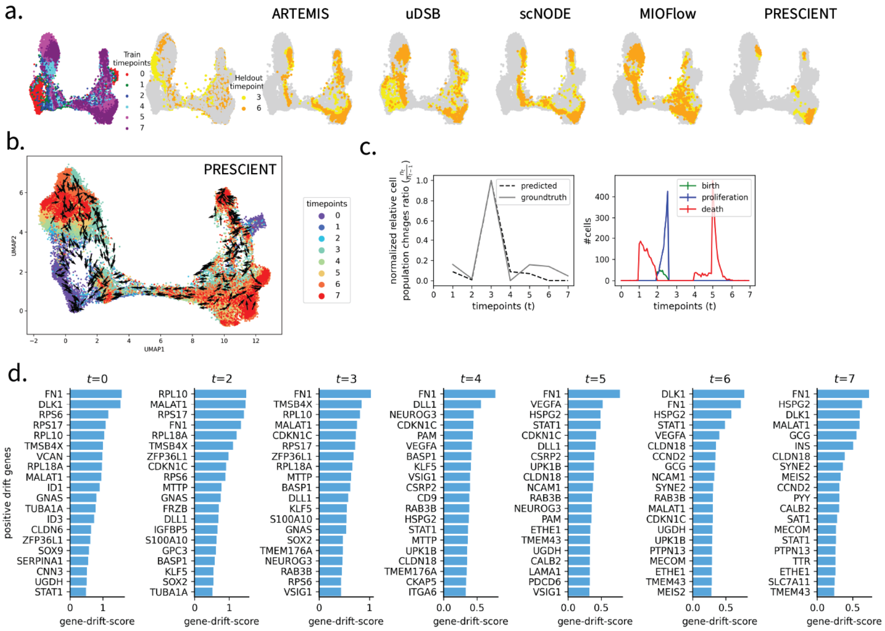


**Supplementary Figure S1.** Application to $\beta$-cell differentiation in human pancreas. a.) Benchmarking ARTEMIS against state-of-the-art methods for predicting gene expression at held-out timepoints (3,6) using pancreatic data, b.) Cell drift inferred by Prescient (w.o. growth rates), c.) Left: Comparison of normalized ratios of relative cell population changes between ground truth and ARTEMIS-predicted cell statuses as live, Right: Number of cells predicted as born, proliferated, and died throughout the trajectory. An increase in cell births and proliferation was observed at $t = 2$ and $t = 3$, coinciding with an increase in relative cell population in the ground truth data. Conversely, a significant number of cells were predicted to die between $t = 1$ to $t = 2$ and $t = 4$ to $t = 6$, aligning with the observed decline in relative cell population, d.)Drift genes identified for zebrafish dataset for remaining ten timepoints.

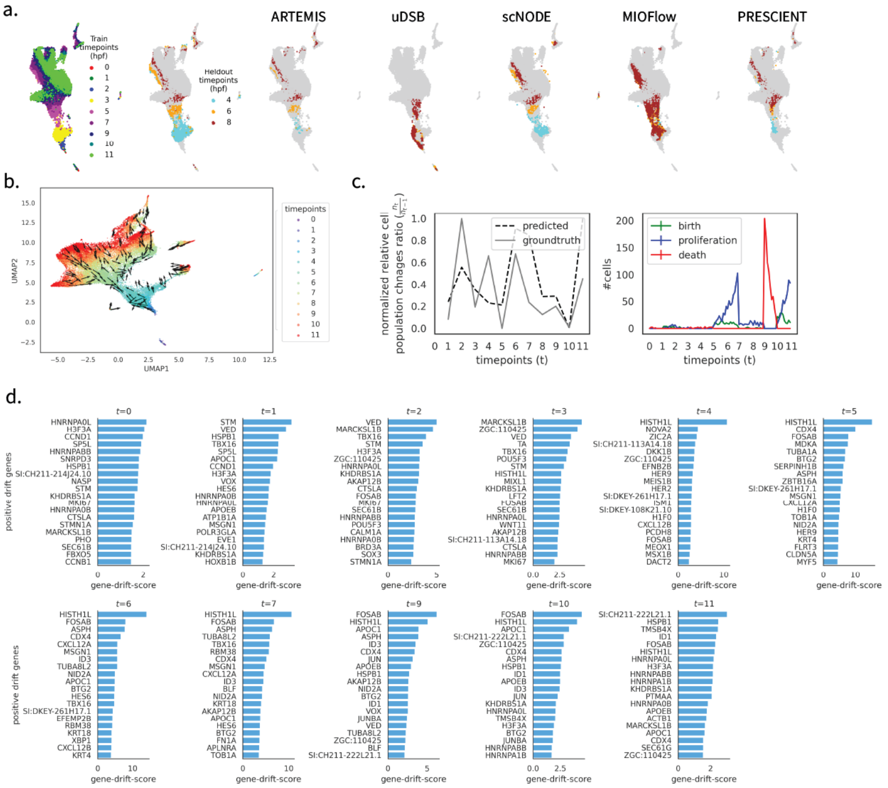


**Supplementary Figure S2.** Application to Zebrafish embryogeneis data, a.) Benchmarking ARTEMIS against state-of-the-art methods for predicting gene expression at held-out timepoints (4, 6, 8) using zebrafish data, b.) Cell drift inferred by PRESCIENT, c.) Left: Comparison of normalized ratios of relative cell population changes between ground truth and ARTEMIS-predicted cell statuses as live, Right: Number of cells predicted as born, proliferated, and died throughout the trajectory. An increase in cell births and proliferation was observed between $t = 4$ to $t = 7$ and $t = 10$ to $t = 11$, coinciding with an increase in relative cell population in the ground truth data. Conversely, a significant number of cells were predicted to die between $t = 9$ to $t = 10$, aligning with the observed decline in relative cell population, d.) Drift genes identified for zebrafish dataset for remaining ten timepoints.

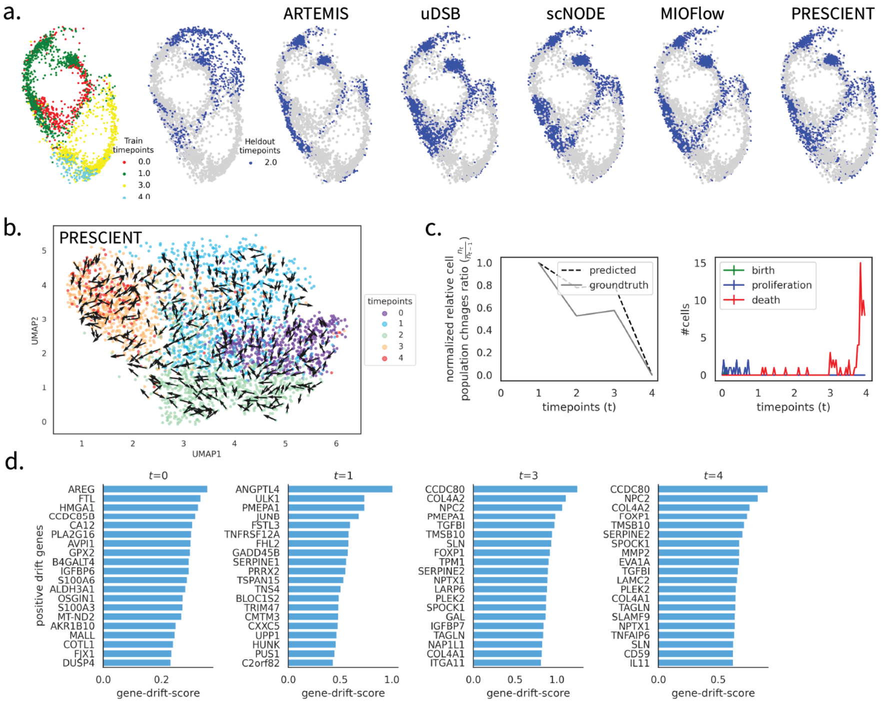


**Supplementary Figure S3.** Application to A549 lung cancer cells treated with *TFGB1* to induce EMT a.) Benchmarking ARTEMIS against state-of-the-art methods for predicting gene expression at held-out timepoint (2) using EMT data, b.) Cell drift inferred by PRESCIENT, c) Left: Comparison of normalized ratios of relative cell population changes between ground truth and ARTEMIS-predicted cell statuses as live, Right: Number of cells predicted as born, proliferated, and died throughout the trajectory. Shorter intervals of predicted cell proliferation are observed earlier in the trajectory, potentially reflecting the higher cell numbers in the ground truth data at these early timepoints. Cell death events are distributed across several shorter intervals throughout the trajectory. A significant spike in predicted cell death is observed towards the end of the trajectory, coinciding with a substantial reduction in the relative cell population in the ground truth data., d.) Drift genes identified for emt dataset for remaining four timepoints.

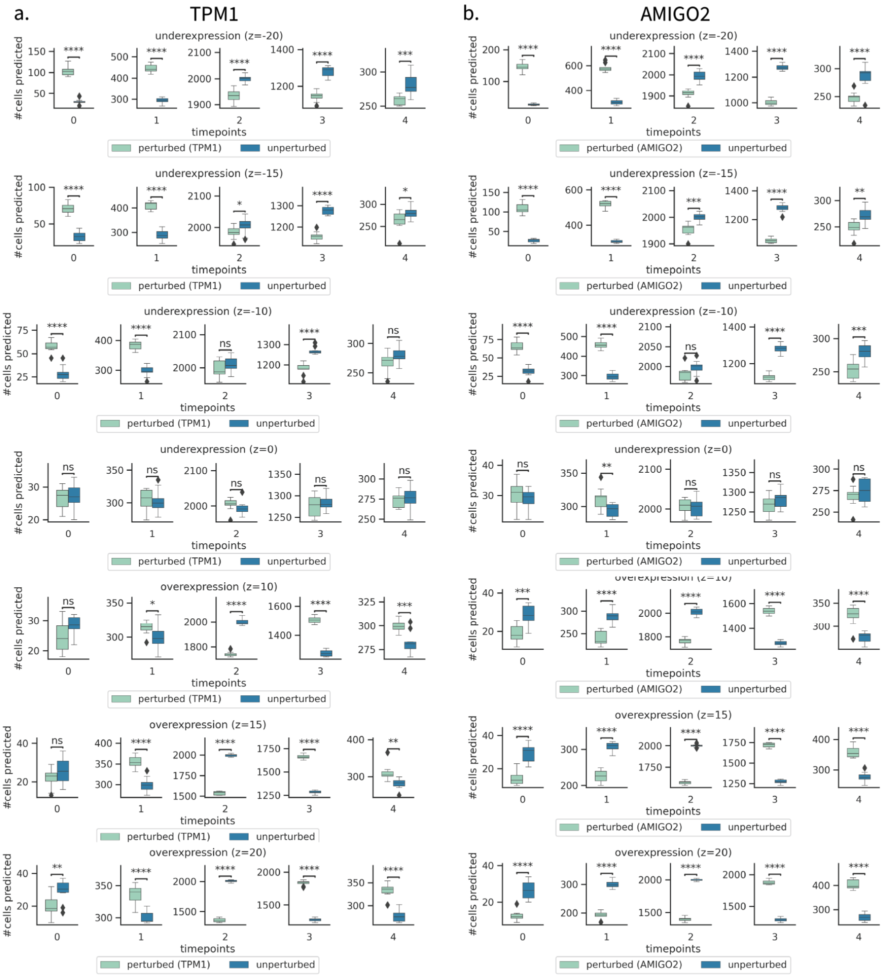


**Supplementary Figure S4.** Perturbation results for different levels of over and underexpression of genes *TPM1* and *AMIGO2*: (-20,-15,-15,-10,0,10,15,20). Cells are assigned to specific timepoints by an MLP classifier, and the number of cells generated from perturbed and unperturbed trajectories were compared using a two-sided t-test at $p < 0.05$.

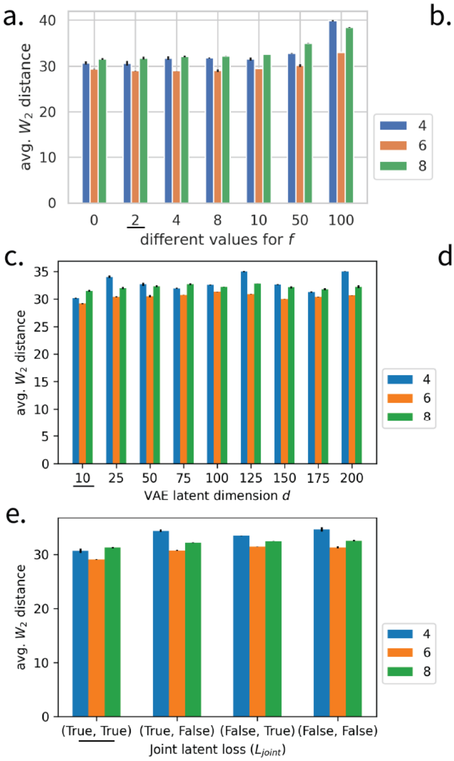


**Supplementary Figure S5.** Evaluating different hyperparameter sets and training configurations for ARTEMIS on the zebrafish dataset [13]. The hyperparameter value underlined is the one used by the baseline model. The different hyperparameters/configurations are: a.) base drift $f$, b.) discretization steps, c.) VAE latent dimension$d$, d.) SDE sampling batch size, e.) Effect of using $L_{joint} = W_2(Z_{\varphi,t}, \overrightarrow{Z}_t) + W_2(X_t, p_\phi(Z_{\varphi,t}))$ loss terms. The x-axis $(\cdot,\cdot)$ indicates when either of the $W_2$ losses is used.
